# Brain activity of professional investors signals future stock performance

**DOI:** 10.1101/2023.05.03.539062

**Authors:** Leonard D. van Brussel, Maarten A.S. Boksem, Roeland C. Dietvorst, Ale Smidts

## Abstract

A major aspiration of investors is to better forecast stock performance. Interestingly, emerging ‘neuroforecasting’ research suggests that brain activity associated with anticipatory reward relates to market behavior and population-wide preferences, including stock price dynamics. In this study we extend these findings to professional investors processing comprehensive real-world information on stock investment options while making predictions of long-term stock performance. Using functional magnetic resonance imaging (fMRI), we sampled investors’ neural responses to investment cases and assessed whether these responses relate to future performance on the stock market. We find that our sample of investors could not successfully predict future market performance of the investment cases, confirming that stated preferences do not predict the market. Stock metrics of the investment cases were not predictive of future stock performance either. However, as investors processed case information, nucleus accumbens (NAcc) activity was higher for investment cases that ended up overperforming in the market. These findings remained robust, even when controlling for stock metrics and investors’ predictions made in the scanner. Cross-validated prediction analysis indicated that NAcc activity could significantly predict future stock performance out-of-sample above chance. Our findings resonate with recent neuroforecasting studies and suggest that brain activity of professional investors may help in forecasting future stock performance.

**Significance Statement:** The investors’ dream of forecasting the stock market is typically considered to be just that: An unrealistic aspiration. However, we find that forecasting stock performance may in fact not be completely unattainable. Results of our neuroimaging experiment reveal that professional investors fail to accurately predict long-term stock performance. However, while processing complex information pertaining to investment cases, brain activity in a region associated with reward anticipation was increased for stocks that would end up overperforming in the future market. Remarkably, this effect held after controlling for the stock information presented in the investment cases. Our findings add to recent work in ‘neuroforecasting’, demonstrating that market behavior can be forecasted by brain activity of a small sample, here of professional investors.

## Introduction

Despite investors’ dedication, consistently forecasting the stock market remains notoriously difficult, if not entirely impossible. Stock markets are inherently governed by unforeseen events, such as political upheaval or natural disasters. Further complications arise from the interplay between sophisticated investors’ preferences and intuitive decisions of naïve actors (1). Accordingly, within the context of traditional finance theory it is generally assumed that it is not possible for investors to reliably forecast the stock market (2) Indeed, while some exceptions have been documented (3, 4), even the most accurate predictions explain, at best, only a relatively small amount of variance in actual stock performance. Still, this does not stop investors from improving their methods. Current improvements seem to result from automation, with stock markets making room for algorithmic trading (5, 6), thereby shifting away from predictions of human investors. Interestingly, however, emerging evidence suggests that neural components of human choice behavior may in fact also help in forecasting market-level behavior.

Recent findings have demonstrated that neural activity associated with individual choice can also be informative of aggregate choice (7). Thus, neural data could potentially be used to forecast the market (i.e., population-wide behavior). In this approach, called ‘neuroforecasting’, neural activity is collected in relatively small samples of 30 to 40 study participants, which is then related to real-world aggregate-level outcomes (8). For instance, neuroforecasting studies have used neural data to forecast album sales in response to music clips (9), box office results in response to movie trailers (10), loan funding rates in response to microloan appeals (11) and crowdfunding proposals (12), advertising elasticity in response to advertisements (13), ad-related click-throughs in response to persuasive messages (14, 15), ad recall of television advertisements (16) and online views for YouTube videos (17).

Neuroforecasting studies sample anticipatory brain activity that occurs before individuals make a conscious choice. Though exceptions exist (15, 16), the majority of neuroforecasting studies focusses on brain activity sampled from three distinct brain areas: predominantly the Nucleus Accumbens (NAcc), or entire Ventral Striatum (VS), the (ventral) Medial Prefrontal Cortex (MPFC), and the Anterior Insula (AIns) (7). It has been proposed that activity in these brain regions offers an opportunity to forecast population behavior because it represents a universal, generalizable response toward the stimulus under investigation (8). Arguably, the stock market reflects collective choice. As such, a generalizable brain response to stock information might be informative of the future performance of stocks as well.

It has indeed been proposed that physiological signals may contain information that can be exploited to better model and predict financial markets (18). Some evidence suggests that a trader’s interoceptive ability as indicated by physiological measures (i.e., heart rate variability) might be informative of success on trading floors (19). Indeed, the brain may contain relevant information for professional investors as well (20). To date, however, relatively little research has investigated brain activity in the context of real-world stock market performance. Previous work has focused on VS activity tracking reactions to corporate earnings news (21), or activation in AIns in response to timely exiting stock bubbles, thereby reaping higher returns (22). Recently, Stallen et al. (23) found initial evidence that anticipatory brain activity measured during assessment of stock prices could forecast their future dynamics. In their study, a sample of university students were shown price graphs of real stocks and were asked to predict whether the price in the next period would go up or down. It was found that average brain activity in the AIns forecasted stock price dynamics, such that higher AIns activity predicted a price inflection, thereby extending the success of neuroforecasting studies to the stock market.

Building on these findings, we aimed to test whether the brain would also be informative of the stock market in a more ecologically valid context, by inviting professional investors to predict stock performance while undergoing fMRI. Specifically, investors were asked to evaluate investment cases of anonymized real-world companies and assess whether the stock of the companies would overperform or underperform in its market segment (benchmark) one year in the future. The stock data of the investment cases was sampled between 2000-2011, and selected such that half of the stocks would underperform whereas the other half would overperform in their market segment one year later. To avoid effects of participants’ expertise in a specific sector, the cases were evenly distributed over multiple sectors. In each sector, both over- and underperforming cases were presented. The investment cases were elaborately specified by providing detailed information pertaining to the financial performance of actual stock profiles, as commonly used by professional investors. This information was presented on five sequentially presented information screens: (1) Company Profile, displaying the sector of the company, its specific industry and its current market capitalization, (2) Price Graph, displaying the performance of the company’s stock in comparison to the sector benchmark, (3) Fundamentals, displaying stock metrics of the past four fiscal years, (4) Relative Valuation, displaying relative valuation information of the company to those of three of its peers, and (5) News Item, displaying a bullet point summary of actual news sampled from a Bloomberg terminal. The duration of the information screens was fixed and differed between screens, based on a pre-test.

We analyzed the fMRI data by extracting neural activation estimates from pre-defined brain volumes of interest (VOIs) which have previously been shown to forecast market-level behavior (8). Specifically, we replicated the approach by Stallen et al. (23) by extracting activity from predefined bilateral foci (8-mm-diameter spheres) in the NAcc, MPFC and AIns. Activation estimates were extracted for each participant, investment case and information screen and then used to inspect brain activity related to stock market performance. Logistic regression analyses were used to test whether brain activity was related to stock market performance, in addition to participants’ predictions and stock metrics. Because stock market performance was identical across participants, we collapsed data at the case level. We thus calculated the average choice that a case would overperform (or not) across participants, and took the average neural activation estimate per information screen per case per VOI. To test the generalizability and replicability of our results we used cross-validation to assess whether brain activity can be used to predict future stock performance.

In line with previous neuroforecasting studies, we hypothesized that the conscious predictions of professional investors would not be predictive of future market performance, whereas their brain activity might. Specifically, we hypothesized that reward-related anticipatory brain activity (i.e., NAcc activity) would forecast whether an investment case would overperform in the future. By contrast, in line with Smith et al. (22), brain activity associated with more general arousal (i.e., AIns activity) might serve as a warning signal, and thus forecast investment cases that would underperform in the future, or price inflection as observed by Stallen et al. (23). Finally, in line with (for example) Falk et al. (14), we hypothesized that brain activity in upstream cortical regions that is related to valuation and subsequent choice behavior (i.e., MPFC), may also be predictive of future performance of the investment cases.

## Materials and Methods

### Participants

Thirty-six professional investors from leading Dutch investment companies took part in the study. On average participants had 19.2 (SD =10.0) years’ experience in the finance industry, 12.4 (SD=9.4) years in asset management and 15.0 (SD=9.6) years in equity analysis. All participants provided written informed consent. Participants received no compensation but were informed that whoever was most accurate in predicting stock outcomes would be awarded a prize of €500. This competition for a prize ensured that participants would predict stock outcome in the task to the best of their abilities. Two participants ended up sharing first place, meaning we evenly split the prize between them, awarding each participant €250. All procedures were conducted as approved by the universities’ ethical Review Board. Exclusion criteria for the study included neurological or cardiovascular diseases, psychiatric disorders, regular drug use, self-reported claustrophobia, or metal parts in the body. Two participants were excluded from the study due to excessive head movement during scanning (i.e., average framewise displacement >0.5mm). A robustness check with all participants included replicated all our main findings. We report data from 34 participants (1 female; mean age 47.6 years, range 29-66, SD=9.1).

### Procedure

Participants were invited to the scanning facility in evening hours. After they had provided informed consent, participants received instructions and completed two practice rounds of the experimental task. Next, participants were placed in the MRI scanner. Structural scans were acquired first, followed by the experiment and functional scanning. Afterwards and outside of the scanner participants provided sociodemographic information, in particular related to their expertise in finance (education, years of financial experience and sectors in which they had most experience). We then asked participants to indicate, for each information screen, how important this information had been for making their predictions (5-point scale), how difficult they found the prediction task and how realistic they found the task (7-point scales). Additionally, we asked them to indicate the fraction of cases they thought they had predicted correctly, and the fraction they thought the other participants had predicted correctly (0-100% in steps of 10%). Next, participants indicated their general willingness to take risks, whether they generally rely on intuition, and whether they in general rely on logic for making decisions (all single items and 10-point response scales). Finally, they were asked to complete questionnaires that assessed thinking-style (rational and experiential; (24)) and open-mindedness (25). Since the focus of the present study is on prediction of future performance of the investment cases (i.e., case-level) and we collapsed all data over participants, differences between individuals (i.e., participant-level) will not be discussed further.

### Stock Performance Prediction Task

To measure brain activity and decision making in response to stock information, we designed the Stock Performance Prediction Task (SPPT). The task was presented using Presentation® software (Version 18.0, Neurobehavioral Systems, Inc., Berkeley, CA, www.neurobs.com). In the SPPT, anonymized investment cases were presented that contained information pertaining to the stock profiles of selected companies. A total of 45 test cases were created by investment experts from a major internationally operating investment company. Due to technical errors with stimulus presentation, for one case one of the information screens was presented incorrectly. As such, we were able to analyze 44 (complete) cases. The investment cases contained actual stock data for three years, all sampled from a period of 11 years (2000 - 2011) to ensure that cases were not affected by a single economic trend. Half of the test cases underperformed its market segment exactly one year later, whereas the other half overperformed. To avoid effects of participants’ expertise in a specific sector, the cases were evenly distributed over nine different sectors: energy, financials, health care, utilities, technology, materials, consumer staples, consumer discretionary and communications. In each sector, both over- and underperforming cases were presented. Investment cases were presented in random order. To prevent any recognition of the cases, participants were not informed about the identities of the investment stocks, or the period from which they were sampled.

Each investment case consisted of five sequentially presented information screens (see: *Stimuli*). Following the five information screens, participants were asked to predict whether the company stock would overperform or underperform in its market segment, one year later. After participants had made their choice, they were asked to rate their confidence in their prediction, choosing between “Quite uncertain”, “Uncertain”, “Certain” and “Quite certain”. Participants did not receive any feedback on their prediction. The case ended with a screen prompting participants to get ready for the next investment case. Every five cases participants were informed of their progress (e.g., ‘Good job, you have completed 5 cases. 40 more to go!’). The total duration of the task was approximately 85 minutes.

### Stimuli

In the SPPT, participants were presented with investment cases consisting of five unique sequential information screens (see Fig. 1 and Table 1. For larger sized stimuli, see *SI Appendix 1*). The first information screen presented an overview of the company (Company Profile screen). Specifically, it showed to which of the nine different market sectors the company belonged, its specific industry and its current market capitalization (in €M). The second information screen presented a performance graph (Price Graph screen), displaying the performance of the company in comparison to the sector benchmark, over the past three years per quartile. The third information screen presented stock fundamentals (Fundamentals screen). Specifically, the fundamentals consisted of sales, earnings before interest and taxes (EBIT), return on equity (ROE) and return on invested capital (ROIC) metrics for the prior fiscal year (FY0) and previous fiscal years (FY-1, FY-2, FY-3). The fourth information screen presented relative valuation information (Relative Valuation screen). This screen compared enterprise value (EV/Sales, EV/EBITDA) and price (P/E, P/B) metrics of the company to those of 3 peers. Finally, the fifth information screen presented a news item (News Item screen). News items consisted of summaries of actual news, sampled from a Bloomberg terminal. Each news item consisted of three bullet points. The first bullet point always indicated the companies’ net income growth over the past year. The second bullet point provided an explanation for this (‘this change was related to…’) and the third item provided a prospect for the future. The duration of each information screen was fixed but differed in length between screens. The duration of the information screens was based on a pre-test such that it assured that participants had enough time to process all information visible on the screen. See Fig. 1 for the respective presentation times per screen.

**Figure 1.**
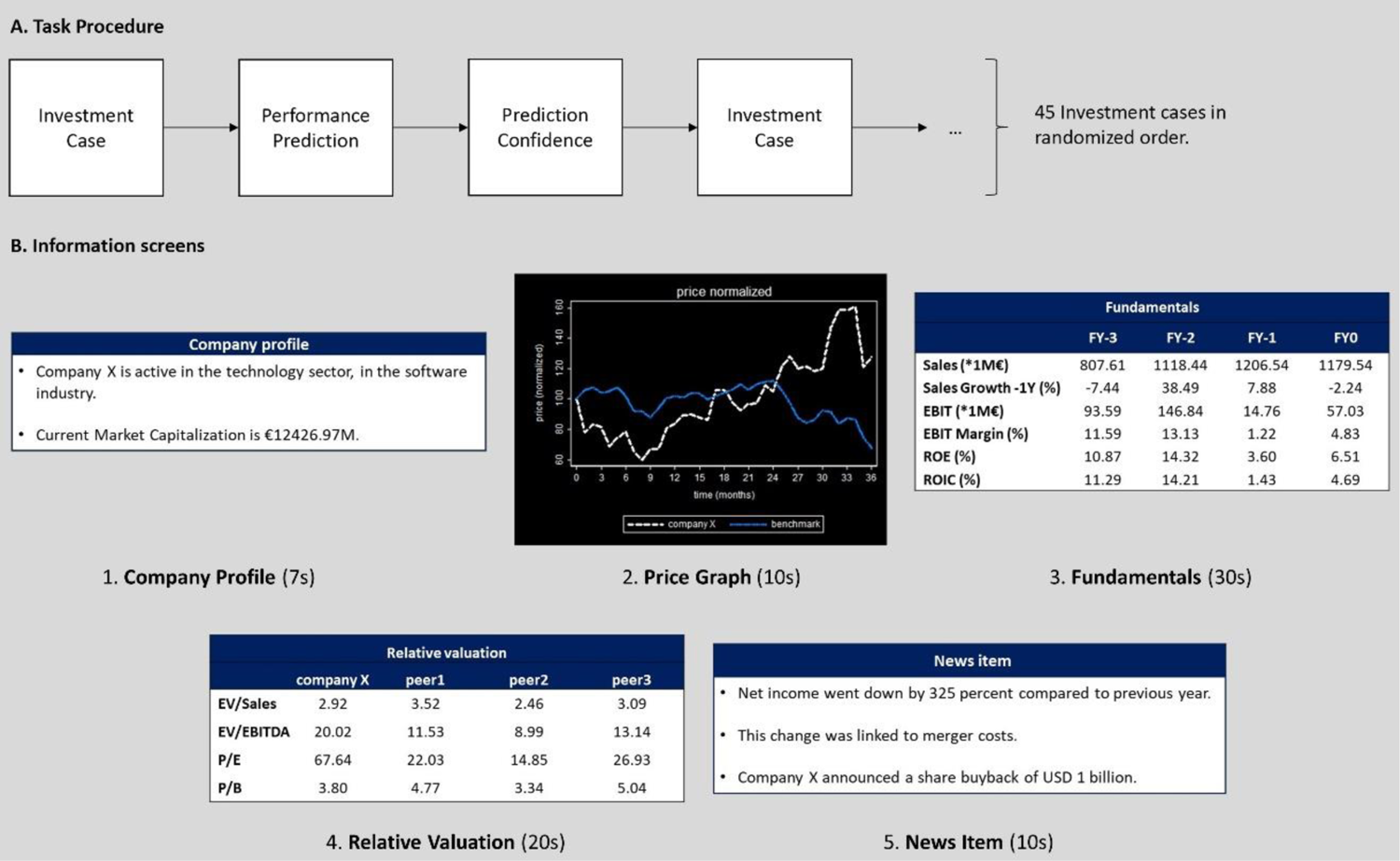
The Stock Performance Prediction Task (SPPT). A. Task procedure: The task consisted of 45 investment cases that were presented in randomized order. Following each investment case, participants were asked to indicate whether they predicted the case to overperform (left button press) or underperform (right button press) in 1 year in the future, as well as their confidence for this prediction. B. Information screens: Each investment case consisted of 5 information screens. From left to right: Company Profile screen (7s), Price Graph screen (10s), Fundamentals screen (20s), Relative Valuation screen (20s) and News Item screen (20s). These screens were presented in sequential order, and jittered. For details on the information presented on the information screens see Table 1.

**Table 1.** Descriptions of the information screens of the SPPT. . Stock metrics in italic were entered as predictors in regression models (all values identical to those presented on the task stimuli, except for CMC which was log-transformed). The Company Profile screen introduced the company (‘Company X’) and indicated its Current Market Capitalization, Industry and Sector. The Price Graph screen presented a price graph, showing the normalized price of Company X with respect to the sector benchmark per quartile for three years. The Fundamentals screen presented stock fundamentals for FY-3, FY-2, FY-1 and FY-0. The Relative valuation screen presented company X valuation with respect to three peers. The News Item screen presented a shortened version of a Bloomberg news item. Each news item began with a statement of the company’s net income growth.

| Screen | Stock metrics | Definition |
| --- | --- | --- |
| Company Profile | *CMC* | Current Market Capitalization. Current shares outstanding / last price. |
|  | *Industry* | Industry of company. |
|  | *Sector* | Sector of company. |
| Price Graph | *P3YRP* | Past 3 Year relative Performance. Assessment of whether the company over- or underperformed w.r.t the sector benchmark over the past 3 years (visually represented). |
| Fundamentals | *Sales* | Sales/revenue/turnover. |
|  | *Sales Growth -1Y (%)* | Sales growth compared to previous year. |
|  | *EBIT* | Earnings before interest and taxes. |
|  | *EBIT Margin (%)* | Earnings before interest and taxes as a percentage of sales. |
|  | *ROE (%)* | Return on Equity. Net income available for common shareholders / average total common equity. |
|  | *ROIC (%)* | Return on Invested Capital. Net operating profit after tax / average invested capital. |
| Relative Valuation | *EV/Sales* | Current enterprise value / trailing 12-month sales. |
|  | *EV/EBITDA* | Current enterprise value / trailing 12-month EBITDA (Earnings Before Interest, Taxes, Depreciation and Amortization). |
|  | *P/E* | Last price / trailing 12-month EPS (Earnings Per Share) before extraordinary items. |
|  | *P/B* | Last price / Book value per share. |
| News Item | *Income growth* | Net percent income growth. |

Between information screens, an inter-trial-interval of a fixation cross with a pseudorandom duration of 2-5s was presented. Following presentation of the last information screen, participants were asked to indicate whether they believed the stock of the company would overperform or underperform the sector benchmark 12 months in the future. Following their prediction, participants were asked to indicate how confident they were of their judgement.

### fMRI Data Acquisition

Imaging was performed with a 3 Tesla MRI scanner (Siemens Verio). Prior to acquisition of functional MRI images, a high-resolution T1-weighted structural image was acquired for anatomical reference (1×1×1 mm, 192 sagittal slices, 9° flip angle). The TE was 30ms and the TR 2300ms. Following acquisition of the structural image, functional scans were acquired by a T2*-weighted gradient-echo, echo-planar pulse sequence in descending interleaved order (3.0 mm slice thickness, 3.0 x 3.0 mm in-plan resolution, 64 x 64 voxels per slice, 90° flip angle). The TE was 25ms, TR 2140ms.

### fMRI Data preprocessing

The fMRI data was first manually inspected to check for anomalies in the data. Next, data of all participants was subjected to MRIqc for quality assessment (26). Inspection of MRIqc’s image quality metrics resulted in the exclusion of two participants (i.e., excessive head-motion, as evidenced by average framewise displacement >0.5 mm). Finally, data of the remaining participants was preprocessed using the standard pipeline of fMRIprep version 20.2.0 (27), based on Nipype (28) (*SI* Appendix 2).

### Data analysis

To test whether brain activity could forecast stock market performance, we first computed whole-brain β-maps using Nibetaseries (29). Rather than looking at changes in raw activity at specific TRs, we chose to look at single-trial activation estimates because these estimates better account for duration differences between the 5 information screens. Specifically, for each participant, we used a least squares-all (LSA) design to obtain a single-trial GLM in which each of the 5 information screens * 44 investment cases were individually modelled with an SPM hemodynamic response function, resulting in a total of 220 trials. Additionally, we included six head-motion regressors, framewise displacement, CSF, WM and global signal as regressors of no interest (all obtained from fMRIprep). This resulted into whole-brain β-maps for each information screen, investment case and participant.

Next, we used a custom Python script to extract activation estimates from predefined volumes of interest (VOIs) from the whole-brain β-maps. We replicated the approach from Stallen et al. (23), by focusing on VOIs which have previously been shown to forecast market-level behavior (8). Specifically, we extracted activity from predefined bilateral foci (8-mm-diameter spheres) in the NAcc (Talairach focus: x, ±10; y, +12; z, −2), the AIns (Talairach focus: x, ±28; y, +18; z, –5), and the MPFC (Talairach focus: x, ±4; y, +45; z, 0). The Talairach coordinates of these VOIs were converted to MNI space, and the average β-estimate for each trial (information screen × investment case) and participant was extracted. The β-estimates were then centered per participant, VOI and information screen. These trial-by-trial activations were then used to inspect brain activity related to stock market performance.

We used logistic regression analyses to test whether brain activity was related to stock market performance, in comparison to stock metrics and participants’ predictions. Because stock market performance was identical across participants, we collapsed data at the case-level, thus averaging brain activity and predictions over participants. We first tested whether participants’ average predictions were related to stock market performance. Next, for each information screen independently, we tested whether neural activity extracted from the three VOIs was related to stock market performance, while controlling for stock metrics (i.e., the key analytical information that was presented to the participants in the experiment; Table 1). Specifically, for the Company Profile screen, we entered the log-transformed current market capitalization (CMC) as stock metric, because CMC was highly skewed. For the Price Graph screen, we entered the past 3 years relative performance (P3YRP) of the company compared to the benchmark at 36 months (i.e., the end of the time period displayed on the graph) as stock metric (P3YRP, 1=overperform, 0=underperform). For the Fundamentals screen, we entered all 5 metrics of the prior fiscal year (FY0). For the Relative Valuation screen, we entered all 4 relative valuation stock metrics of the company. Finally, for the News information screen we included net income growth as stock metric. For the neural activity we took the average β-estimate per information screen per case per VOI (N=44 observations). Additionally, we investigated whether neural activity at the information screens was related to stock performance inflection and participants’ predictions. Stock performance inflection was assessed by taking the P3YRP (above or below the sector benchmark, as indicated on the Price Graph screen) and stock performance one year in the future and establishing whether the direction is identical or not (0 = no inflection, 1 = performance inflection) Thus, stock performance inflection was a binary variable meaning we used identical logistic regression models.

Following tests per information screen, we then proceeded to test whether neural activity was related to stock market performance, controlling for all stock metrics and choice behavior. We first tested whether the stock metrics were related to stock market performance (Market model). Next, we added participants’ predictions to the model (Market + Behavior model). Finally, we added neural activity to the model (Market + Behavior + Brain). The Kaiser-Meyer-Olkin measure indicated that NAcc activity was factorable (KMO=.74). Oblique rotation factor analysis revealed that NAcc activity across the five information screens could be adequately described by two factors explaining 63.0% of variance (Test of the hypothesis that 2 factors are sufficient: χ²(1) =.520, *p* =.472; correlation of the two factors: r= .459). For the Market + Behavior + Brain model, we included the first factor score in the regression model. Regression analyses were performed with custom R code, in combination with the lmertest package 3.1-3 for linear models (30) and the Jtools package 2.1.4 for outputting regression tables (31). For ease of interpretation, all continuous predictors were standardized.

Having identified significant predictors with our regression analyses, we finally tested how well our models could predict market performance. For this, we utilized the caret package (32) to subject our logistic regression models to cross-validation. Specifically, we choose k-fold cross-validation, as recommended by Poldrack et al. (33), setting the test size to 20%, as recommended by Varoquax et al. (34). Given the small sample (N=44) we used stratification to ensure an equal distribution of overperforming and underperforming cases in our test set. As a control, we also performed leave-one-out cross-validation. The significance of resulting accuracies was assessed by exact binomial tests against chance.

## Results

### Investor’s predictions of future stock performance

We found considerable variation in the predictions made by the investors. Over all cases, on average participants predicted 49.5% overperforming (SD = 24.3%). Similarly, we found variation in how well participants could predict the future performance of the investment cases (mean accuracy = 52.6%, SD=24.2%). Overall, participants found the forecasting task difficult (M = 4.41, SD=0.99) but realistic (M = 4.05, SD=1.41), significantly above midpoint of the 7-point scales (both p’s <0.05). When asked for general feedback at the end of the experiment, none of the participants reported to have recognized one or more of the investment cases.

Next, we investigated whether participants’ average prediction of market performance was related to actual performance of the companies one year into the future. A logistic regression model on the 44 cases indicated that participants’ predictions were not significantly related to market performance (*b* =0.222, SE = 0.308, *p*=0.471). This finding is consistent with the efficient market hypothesis (2). We also looked at participants’ self-reported confidence and found no correlation between how well participants could predict the future performance of the investment cases and the average confidence rating of participants’ predictions (*r*=0.02, *p*=0.88).

Finally, we computed the average of participants self-reported importance of the five information screens rated on a 5-point scale from not at all (1) to very important (5). We found that the average self-reported importance of the Company Profile screen was 1.88 (SD = 1.23), the Price Graph screen 2.91 (SD = 0.97), the Fundamentals screen 3.97 (SD = 1.03), the Relative Valuation screen 2.97 (SD=1.19) and the News Item screen 3.91 (SD = 1.29). Post hoc analysis, consisting of multiple pairwise t-tests with p-values adjusted using Bonferroni correction, revealed that participants found the fundamentals screen and news item screen most important for their predictions, and the profile screen least important (all p’s <.05).

### Investor’s brain activity related to future stock performance

Having confirmed that our sample of professional investors could not predict future market performance of the stocks, we next tested our critical hypothesis that brain activity was related to market performance. Further logistic regression analyses investigated neural activity in our 4 VOIs, independently for every information screen, controlling for the stock metrics (Table 2). We found that only average NAcc activity was positively related to stock benchmark overperformance one year in the future, at the Company Profile screen (*b* =1.35, SE = 0.61, *p*=0.029), the Price Graph screen (*b* =1.90, SE = 0.82, *p*=0.020), and the Fundamentals screen, (*b* =1.34, SE = 0.68, *p*=0.047). Across all screens, none of the other neural predictors (MPFC and AIns) or stock metrics were found to be predictive. In reduced models with only NAcc predictors (i.e., without MPFC and AIns) and stock metrics for each screen, neural activity remained significant for the Company Profile screen (*b* =0.98, SE = 0.38, *p*=0.010) and Price Graph screen (*b* =1.17, SE = 0.46, *p*=0.011, but not the Fundamentals screen (*b* =0.47, SE = 0.35, *p*=0.186). To further illustrate our findings, we conducted two-sample t-tests to compare NAcc activity for underperforming cases vs. overperforming cases. We found that average NAcc activity for underperforming cases was lower than for overperforming cases at the Company Profile screen (t_(42)_ = −2.92, p=0.005) and Price Graph screen (t_(42)_ = −2.65, p=0.011), but not for the other information screens (Fig. 2).

**Figure 2.**
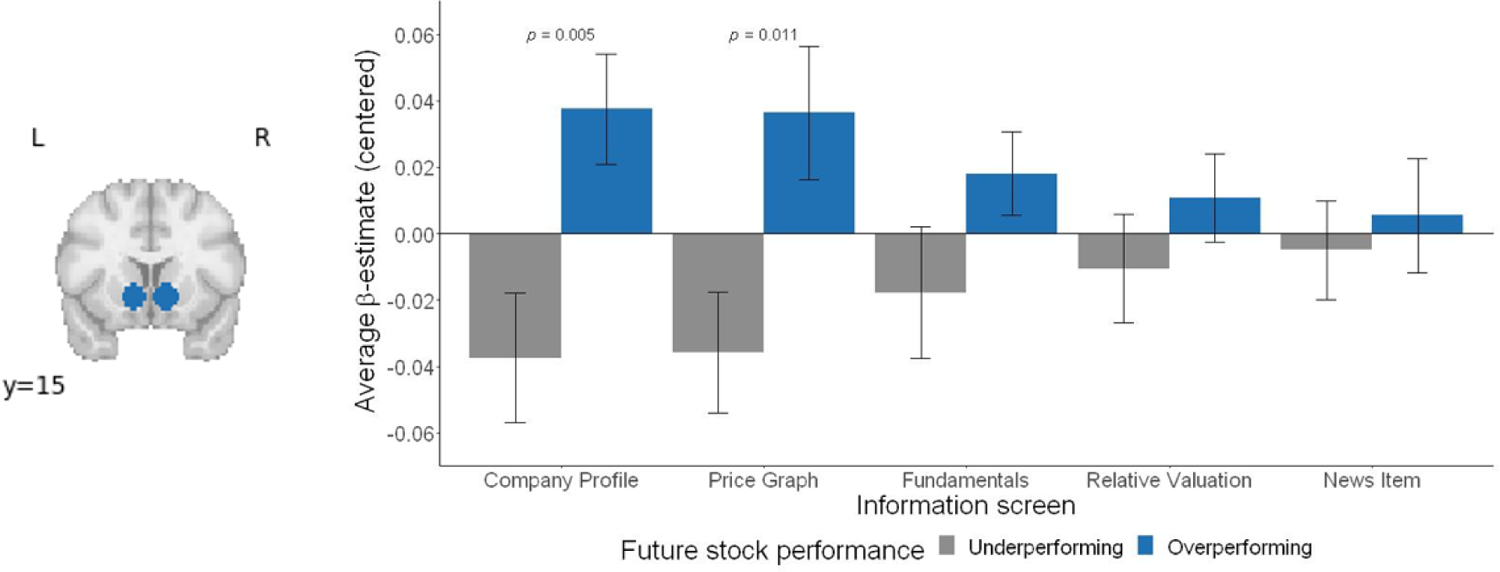
Nucleus Accumbens activity relates to future stock performance. Left: NAcc VOIs. Right: NAcc activity for the five information screens. NAcc activity is significantly higher at the company profile screen and price graph screen for overperforming cases vs. underperforming cases (1 year in the future). Average NAcc activity for overperforming cases was higher than for underperforming cases at the Company Profile screen (t_(42)_ = 2.92, p=0.005) and Price Graph screen (t_(42)_ = 2.65, p=0.011), but not for the other information screens (all p’s >0.05).

**Table 2.** Logistic regression models of neural activity and stock metrics, forecasting future market performance (overperformance (1) vs. underperformance (0)) of company stocks, analyzed per information screen.

|  | Company Profile | Price Graph | Fundamentals | Relative Valuation | News Item |
| --- | --- | --- | --- | --- | --- |
| Current Market Capitalization | 0.30 (0.35) |  |  |  |  |
| Profile - NAcc | **1.33 \* (0.61)** |  |  |  |  |
| Profile - MPFC | 0.32 (0.60) |  |  |  |  |
| Profile - AIns | -0.75 (0.55) |  |  |  |  |
| P3YRP |  | 1.00 (0.84) |  |  |  |
| Graph - NAcc |  | **1.90 \* (0.82)** |  |  |  |
| Graph - MPFC |  | -0.75 (0.57) |  |  |  |
| Graph - AIns |  | -0.17 (0.74) |  |  |  |
| Sales |  |  | -0.27 (0.46) |  |  |
| EBIT |  |  | 0.05 (0.43) |  |  |
| EBIT Margin (%) |  |  | 0.27 (0.49) |  |  |
| ROE (%) |  |  | -0.28 (0.73) |  |  |
| ROIC (%) |  |  | -0.17 (0.59) |  |  |
| Fundamentals - NAcc |  |  | **1.34 \* (0.68)** |  |  |
| Fundamentals - MPFC |  |  | 0.09 (0.51) |  |  |
| Fundamentals - AIns |  |  | -1.11 (0.75) |  |  |
| EV/Sales |  |  |  | 0.25 (0.64) |  |
| EV/EBITDA |  |  |  | 1.06 (0.69) |  |
| P/E |  |  |  | -0.83 (0.69) |  |
| P/B |  |  |  | 0.78 (1.92) |  |
| Relative Valuation - NAcc |  |  |  | 0.54 (0.57) |  |
| Relative Valuation - MPFC |  |  |  | 0.43 (0.50) |  |
| Relative Valuation - AIns |  |  |  | -0.78 (0.57) |  |
| Income growth |  |  |  |  | 1.17 (1.48) |
| News item - NAcc |  |  |  |  | -0.32 (0.50) |
| News item - MPFC |  |  |  |  | 0.24 (0.50) |
| News item - AIns |  |  |  |  | 0.40 (0.50) |
| N | 44 | 44 | 44 | 44 | 44 |
| AIC | 60.53 | 59.62 | 72.76 | 69.07 | 66.98 |
| Pseudo R2 (McFadden) | 0.17 | 0.19 | 0.10 | 0.13 | 0.07 |

Statistics are coefficients with SEMs in parentheses. All continuous predictors are mean-centered and scaled by 1 standard deviation. \* $p < 0.05$ .

Additionally, we tested whether brain activity was related to stock performance inflection. Stock performance inflection is defined here as a change in the direction of the stock performance between the current stock performance (above or below the benchmark as indicated on the Price Graph screen) and performance 1 year in the future. For stock performance inflection, we again used logistic regression (0 = no inflection, 1 = performance inflection) to investigate neural activity independently for every information screen. We found that none of the neural predictors (NAcc, AIns and MPFC) were related to stock performance inflection, for any of the information screens (all p’s > 0.05).

Next, we investigated whether NAcc activity was related to future stock performance in combination with participants’ prediction (behavior) and stock metrics (market). We observed that NAcc activity across the five information screens was positively correlated (NAcc median: *r* = 0.50). Oblique rotation factor analysis revealed that activity across the five screens could be sufficiently described by two factors (See Materials and Methods). The first factor combines NAcc activity at the Company Profile screen, the Price Graph screen, and the Fundamentals screen, whereas the second factor combines activity of the Relative Valuation, and News Item screen. We found that the first factor was significantly related to market performance (*b* =0.90, SE = 0.39, *p*=0.022), but the second factor was not (*b* =-0.12, SE = 0.32, *p*=0.720). For the full model (i.e., Market + Behavior + Brain) we included only the first factor score of NAcc activity. We then used logistic regression analyses to investigate whether NAcc combined activity would still predict future stock beyond stock metrics (Market) and participants’ prediction (Behavior) (Table 3). We found that overall, the Market model and Market + Behavior model were insignificant (Market model: χ²(12)= 13.099, *p* = 0.362; Market + Behavior model: χ²(13) = 14.440, *p* = 0.344). However, the Market + Behavior + Brain model was significant (χ²(14) = 24.945, *p* = 0.035), with NAcc activity at the Company Profile, Price Graph and Fundamentals screen (i.e., the first factor) as a significant predictor (*b* =1.90, SE = 0.77, *p*=0.013), as well as the P/E stock metric (*b* =-1.91, SE = 0.93, *p*=0.039). However, unlike the first NAcc factor (see above) the P/E stock metric was not significantly related to market performance by itself (*b* = −0.05, SE = 0.31, *p*=0.866). Direct model comparisons indicated that the Market + Behavior + Brain model predicted future stock market performance significantly better than the Market model (F(2,29) = 11.85, *p* = 0.003) and the Market + Behavior model (F(2,29) = 10.50, *p* = 0.001).

**Table 3.** Comparison of Market, Market + Behavior and Market + Behavior + Brain logistic regression models forecasting future market performance (overperformance (1) vs. underperformance (0)) of company stocks.

|  | Market | Market + Behavior | Market + Behavior + Brain |
| --- | --- | --- | --- |
| Current Market Capitalization | 0.75 (0.77) | 0.86 (0.76) | 1.20 (1.08) |
| P3YRP | 0.19 (0.85) | -0.07 (0.90) | 0.45 (1.09) |
| Sales | -0.36 (0.48) | -0.20 (0.50) | 0.95 (0.75) |
| EBIT | -0.29 (0.58) | -0.58 (0.62) | -1.36 (0.91) |
| EBIT Margin (%) | -0.15 (0.73) | -0.03 (0.80) | 0.45 (1.11) |
| ROE (%) | -0.41 (0.94) | -0.37 (0.96) | -1.14 (1.58) |
| ROIC (%) | -1.01 (0.83) | -1.30 (0.97) | -1.09 (1.14) |
| EV/Sales | 0.00 (0.89) | 0.05 (0.89) | 1.06 (1.02) |
| EV/EBITDA | 1.99 (1.21) | 1.98 (1.24) | 2.56 (1.48) |
| P/E | -1.22 (0.77) | -1.26 (0.72) | **-1.91 \* (0.93)** |
| P/B | 1.25 (3.68) | 1.03 (3.30) | 0.21 (1.13) |
| Income growth | 1.27 (1.52) | 0.55 (1.57) | 2.33 (1.87) |
| Prediction |  | 0.57 (0.50) | 1.31 (0.77) |
| NAcc – Company Profile,<br>Price Graph, Fundamentals |  |  | **1.90 \* (0.77)** |
| N | 44 | 44 | 44 |
| AIC | 73.90 | 74.56 | 66.05 |
| Pseudo R2 (McFadden) | 0.21 | 0.24 | 0.41 |

Statistics are coefficients with SEMs in parentheses. All continuous predictors are mean-centered and scaled by 1 standard deviation. \* $p < 0.05$ .

### Predicting future stock performance

Finally, we were interested to see how well we could predict future stock market performance. To test this, we applied stratified 5-fold cross-validation to logistic regression models of stock metrics (Market), participants’ predictions (Behavior) and the first factor score of NAcc activity (Brain). A model trained on stock metrics predicted future case performance with 43.18% accuracy, 95% CI [28.35, 58.97], not exceeding chance (*p* = 0.854, exact binomial test). Likewise, a model trained on investors’ predictions had an accuracy of 43.18%, 95% CI [28.35, 58.97], not exceeding chance either (*p*=0.854, exact binomial test). However, a model trained on the first factor score of NAcc activity predicted future case performance with 68.18% accuracy, 95% CI [52.42,81.39], exceeding chance (*p*=0.011, exact binomial test). As a control, we also tested prediction accuracies for our models using a leave-one-out cross-validation approach. Here too, we found that accuracy of the model trained on the first factor score of NAcc activity exceeded chance (accuracy = 65.91%, *p*=0.024), but that the model trained on stock metrics (accuracy =43.18%, *p*=0.854) or participants’ predictions (accuracy = 50.00%, *p*=0.560) did not.

## Discussion

We investigated whether brain responses of professional investors to complex real-world investment cases relate to future stock performance. We found that investors’ brain activity during exposure to stock information is indicative of population-wide investment decisions (i.e., aggregate choice). By contrast, their collective predictions (i.e., group choice behavior) did not forecast future stock market performance, consistent with the efficient market hypothesis (2). Our results extend previous findings in neuroforecasting stock markets (23) by sampling brain activity from a select group of professionals, with high-level expertise in making investment decisions. While several studies have shown that NAcc activity from a small sample can be used to predict market-level outcomes (7), these studies typically recruited university students as participants. Moreover, most neuroforecasting studies to date have dealt with relatively simple stimuli that had to be evaluated in only a few seconds (e.g., crowdfunding options described in only a few sentences; (12)). Our study shows that NAcc activity also seems predictive when professionals are extensively processing and evaluating complex information, that untrained eyes would struggle to interpret.

More specifically, we found that NAcc activity at the Company Profile screen specifying the company’s sector, industry, and market capitalization and NAcc activity at the Price Graph screen showing the stock’s performance in comparison to the sector benchmark over a three-year period, relates to future stock performance. At the Fundamentals screen, displaying stock metrics of the past four years, this forecasting relationship was reduced but still significant in combined models. NAcc activity, being associated with positive valuation and anticipated reward (35), has previously been found to respond to corporate earnings (21) and the tracking of the magnitude of a price bubble (22). In a single-patient case study, NAcc dopamine release was also found to track the market price in an experimental sequential-investment task (36). Our results suggest that in our experiment, investors particularly assessed the expected value of the company stocks based on information presented early on, generating a brain response related to future stock performance.

In our experimental design we presented the information screens always in the same order. As such, we cannot determine whether the neural response was elicited by the specific information presented at the Company Profile screen and Price Graph screen, or because this was the very first information that was presented to the professional investors. Other neuroforecasting studies have reported initial NAcc activity, occurring at the very first moments of stimulus presentation, to be most predictive of market-level success (11, 12). Relative to the entire duration of each investment case (77 seconds, excluding inter-trial intervals between information screens), the Company Profile screen (7 seconds) and Price Graph screen (10 seconds) comprise only the initial phase of presentation of information of each investment case, giving a first indication of its characteristics. During presentation of subsequent information (i.e., the Fundamentals, Relative Valuation, and News Item screens), the initial anticipatory response may wash out. A speculative explanation is that over time, investors become more involved with integrating all information and making a deliberate choice, which may diminish the impact of their first intuitive response to the case. This resonates with new theories in behavioral economics, proposing that initial valuation in the decision-making process can become obscured by cognitive noise, caused by complexity of the task at hand (37, 38). On a neural level, previous work hints at involvement of MPFC in this process (39, 40). However, we find no relationship between MPFC activity and future stock performance, which is in line with the findings of Stallen et al. (23).

We also find that AIns activity is not predictive of future stock performance. AIns activity has been associated with negative or generally aroused affect, as well as avoidance behavior (41, 42), and may serve as a warning signal in financial trading (22). Indeed, Stallen et al. (23) found that AIns activity was related to stock price inflections in the next period, thus reflecting changing demand for stocks and associated price decreases. In our study we did not find AIns activity to be related to price inflection. However, we could only assess stock performance inflection by comparing differences in market performance over the period of one year. Moreover, in the experiment of Stallen et al. (23) participants made 10 consecutive investment choices per stock (each for the following day), whereas in our experiment each investment case was assessed only once. It might be that in a context of financial trading, AIns mostly responds to sudden changes of the presented information, which we did not investigate in our experiment. Whether or not this implies that AIns responses can only be related to short-term market outcomes, or also long-term, remains an open question. The alleged different mechanisms of the brain components investigated here (i.e., NAcc, MPFC, AIns), and their potential for neuroforecasting, deserves more attention (7, 8) and further stresses the importance of studying the neurobiology of financial decision-making (20).

Though the objective stock information presented to the participants resembles typical input of trading algorithms (5, 6), our results found no relationship between stock metrics and future stock performance, except for the Price-to-Earnings ratio (P/E stock metric, Relative Valuation screen) which was only significantly related to future market performance in the combined model (Market + Behavior + Brain). Instead, we found brain activity in response to the stock metrics to be informative of future market performance. This finding begs the question how the brain of the professional investors processed information that by itself was unrelated to future stock market performance into a signal capable of predicting the future market. While in our model the stock’s Current Market Capitalization (CMC; Company Profile screen) and Past 3 Year Relative Performance (P3YRP; Price Graph screen) were unrelated to future market performance and investors’ perceived importance of the information screens was also lowest for these two screens, NAcc activity during these information screens did predict future stock performance. We speculate that the financial expertise and extensive experience of the professional investors enabled them to combine and integrate the provided information with their prior knowledge of similar investment cases, resulting into an intuitive response. This anticipatory, intuitive response appears to be of great value, as our fMRI data suggests. This dovetails with findings that financial traders’ intuition affects their performance (43, 44, 20). Where some accounts emphasize cognitive reflection and pattern recognition as key mechanisms for trader intuition (43, 44), more recently it has been argued that trader intuition might be more emotional (20). In our experiment, pattern recognition of the presented information may have elicited an anticipatory (and possibly affective) NAcc response. The extent to which investors are conscious of such (positive) anticipation, and whether and how this response is modulated by their expertise and experience remain interesting questions for future studies.

Our study advances neuroforecasting literature by presenting complex, realistic real-world investment information to professional investors predicting stock performance one year in the future. In line with the results of Stallen et al. (23), we find that anticipatory brain activity forecasts future market performance, while behavior or stock metrics do not. Our study thus offers a conceptual replication of the predictive role of brain activity in financial markets with increased ecological validity. Although the nature of our data necessitates that our results were obtained in a lab-environment, we exposed our participants to realistic investment scenarios mimicking financial decision-making in real-world institutions (45). In fact, the information that we presented would be mostly unintelligible to untrained eyes, underlining the importance of our sample of professional investors with specific and relevant expertise. While previous studies found a relationship between market-level outcomes and brain responses of naïve participants to simplified stimuli presented out-of-context (7), our study shows that the market can be predicted by brain responses of experts processing comprehensive, realistic information.

There are also limitations to our work. The complexity of the investment cases that we presented allowed for a relatively low number of stocks that we could present (n = 44), while still having a relatively long experimental duration. In addition, even though we anonymized the historical investment cases and none of the investors reported that they recognized a case, we cannot fully rule out the possibility of (unconscious) recognition of some anonymized cases, which might have contributed to the neural responses. Finally, because information screens of the investment cases were presented sequentially and always in the same order, we cannot draw any conclusions on how the information exactly influenced brain activity. This generates several interesting questions for future research, for example whether specific financial information drives brain activity predictive of future stock performance, or whether it is the initial response specifically that is predictive of market performance, independent of the exact type of information.

In conclusion, our study suggests that future stock market performance can be predicted by brain measures of professional investors. This challenges traditional theoretical accounts (2), and raises the question whether financial institutions should invest in collecting such information. However, we acknowledge the exploratory nature of our findings and believe that it is too early to suggest that neural measures should become an integral component of investment institutions. More evidence is needed to thoroughly understand the specific role of neural components underlying investment decisions and their potential for neuroforecasting. In particular, evidence should be gained from prospective neuroforecasting studies (as opposed to retrospectively predicting ‘future’ market performance). On the other hand, together with the findings of Stallen et al. (23) and Smith et al. (22) and general neuroforecasting research (7), accumulating evidence does suggest that humans, including professional investors, may share a neural response to stimuli that is related to future market-level performance, suggesting to financial institutions the investment value of collecting such information.

## SI Appendix 1 Example information screens of the Stock Performance Prediction Task

To provide a better understanding of the information screens that participants were shown in the Stock Performance Prediction Task (SPPT), for each of the 5 information screens, two examples are shown below. Note that for all information screens, the example on the top corresponds to the example information screen depicted in Fig. 1.

**Fig. S1.**
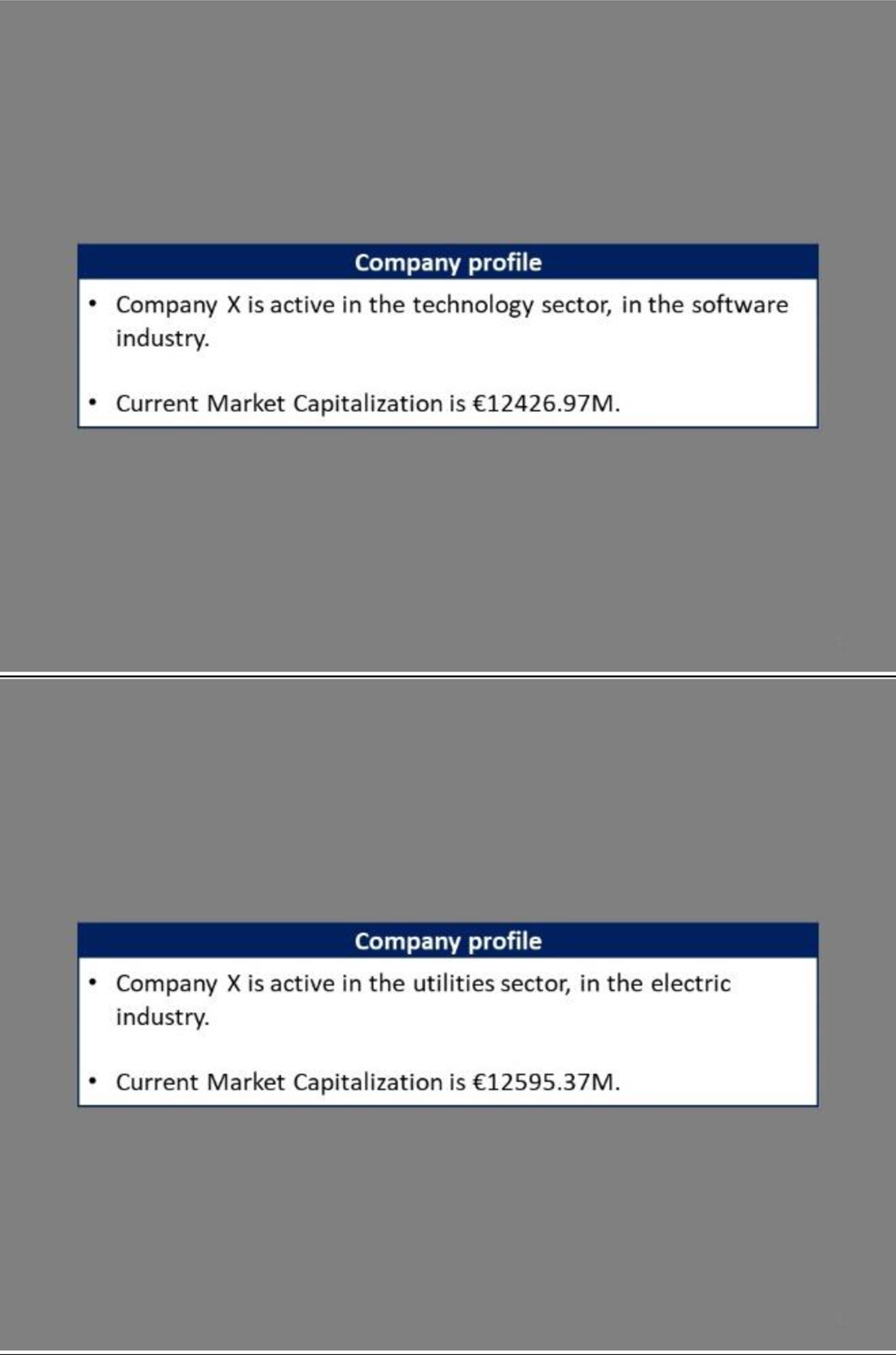
Two examples of the Company Profile screen. The Company Profile screen showed which of the nine different market sectors the company belonged to, its specific industry and its current market capitalization (in €M).

**Fig. S2.**
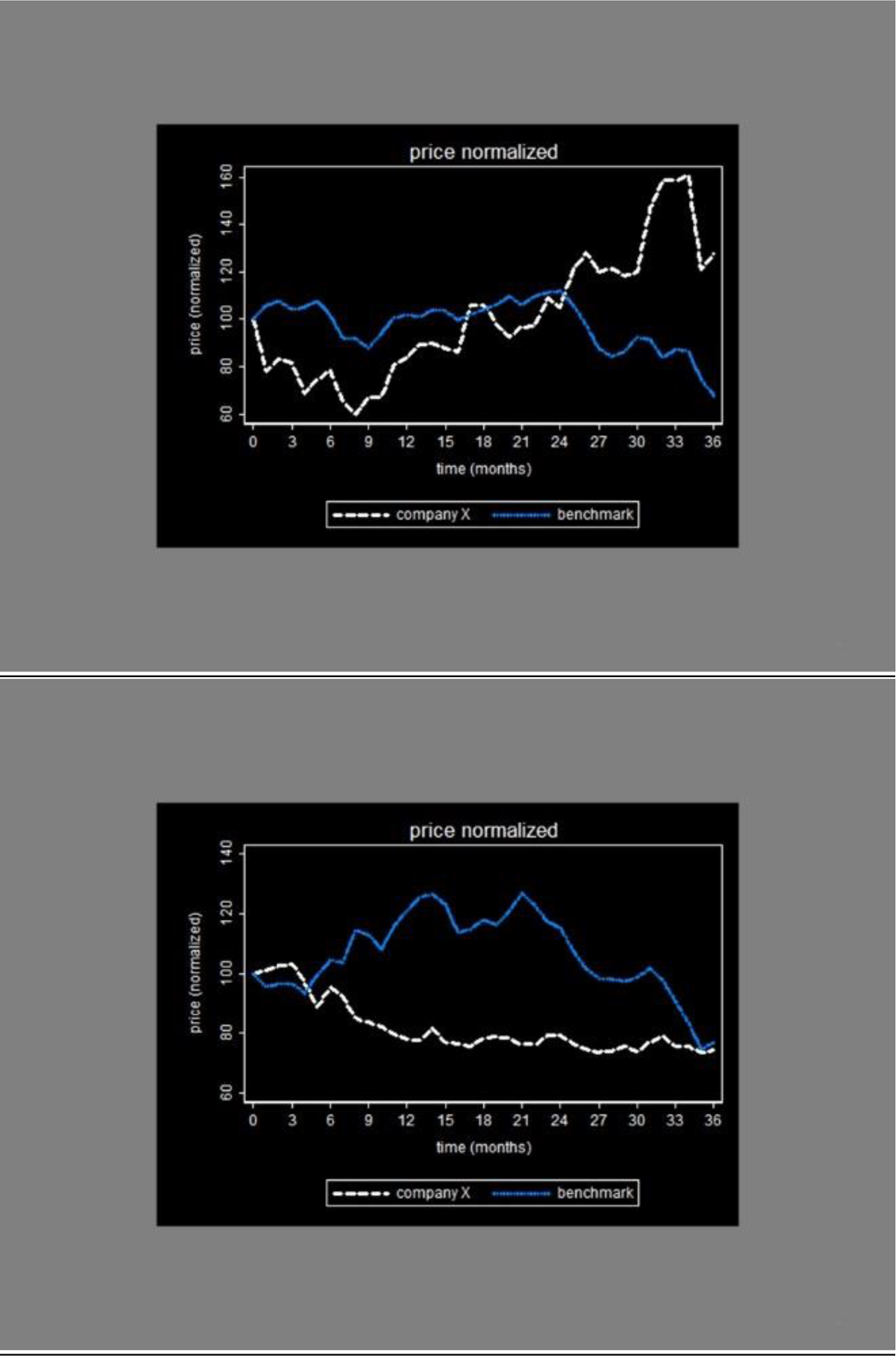
Two examples of the Price Graph screen. The Price Graph screen displayed the performance of the company in comparison to the sector benchmark, over the past three years per quartile.

**Fig. S3.**
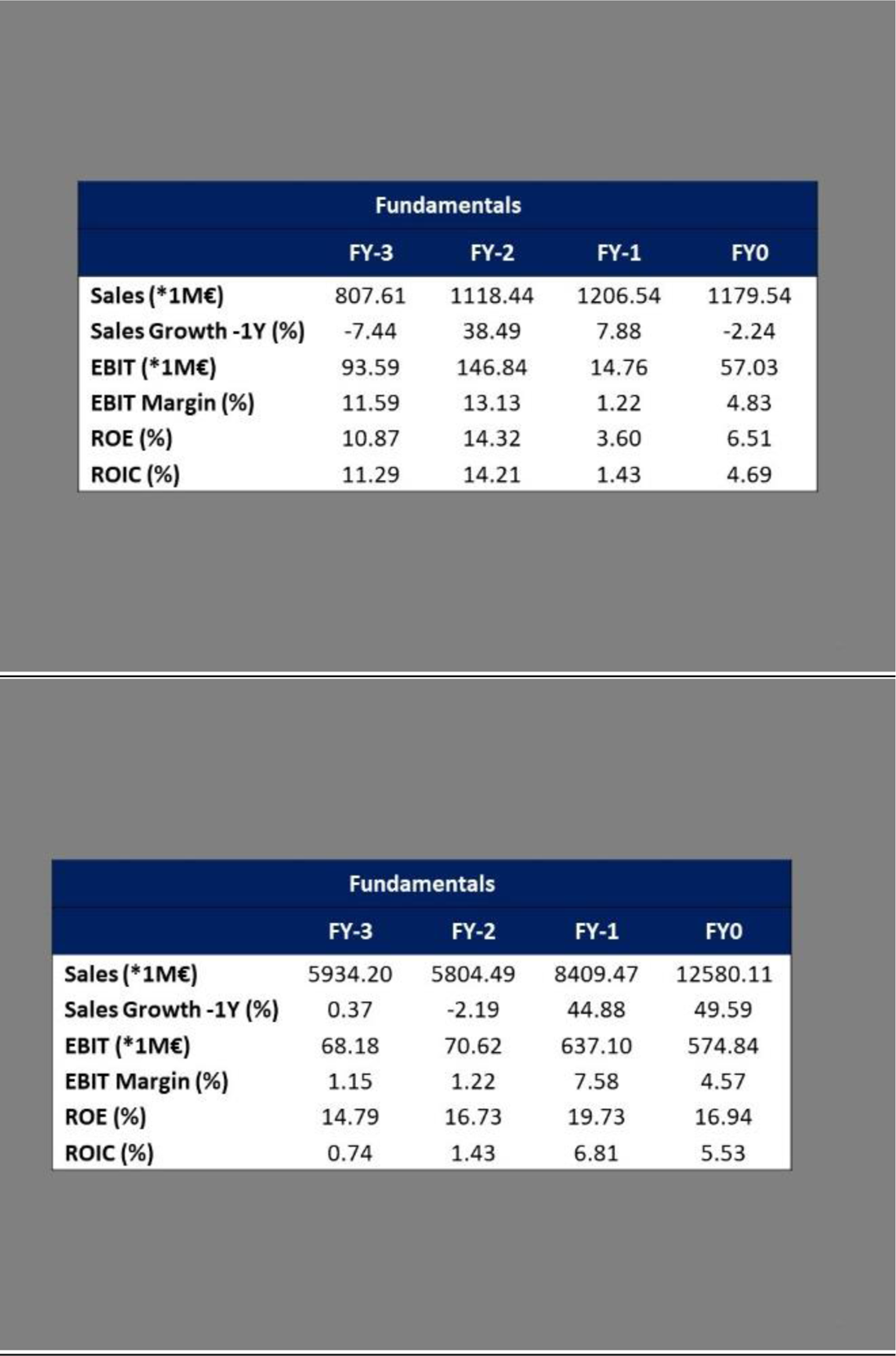
Two examples of the Fundamentals screen. The Fundamentals screen presented stock fundamentals, consisting of sales, earnings before interest and taxes (EBIT), return on equity (ROE) and return on invested capital (ROIC) metrics for the prior fiscal year (FY0) and previous fiscal years (FY-1, FY-2, FY-3).

**Fig. S4.**
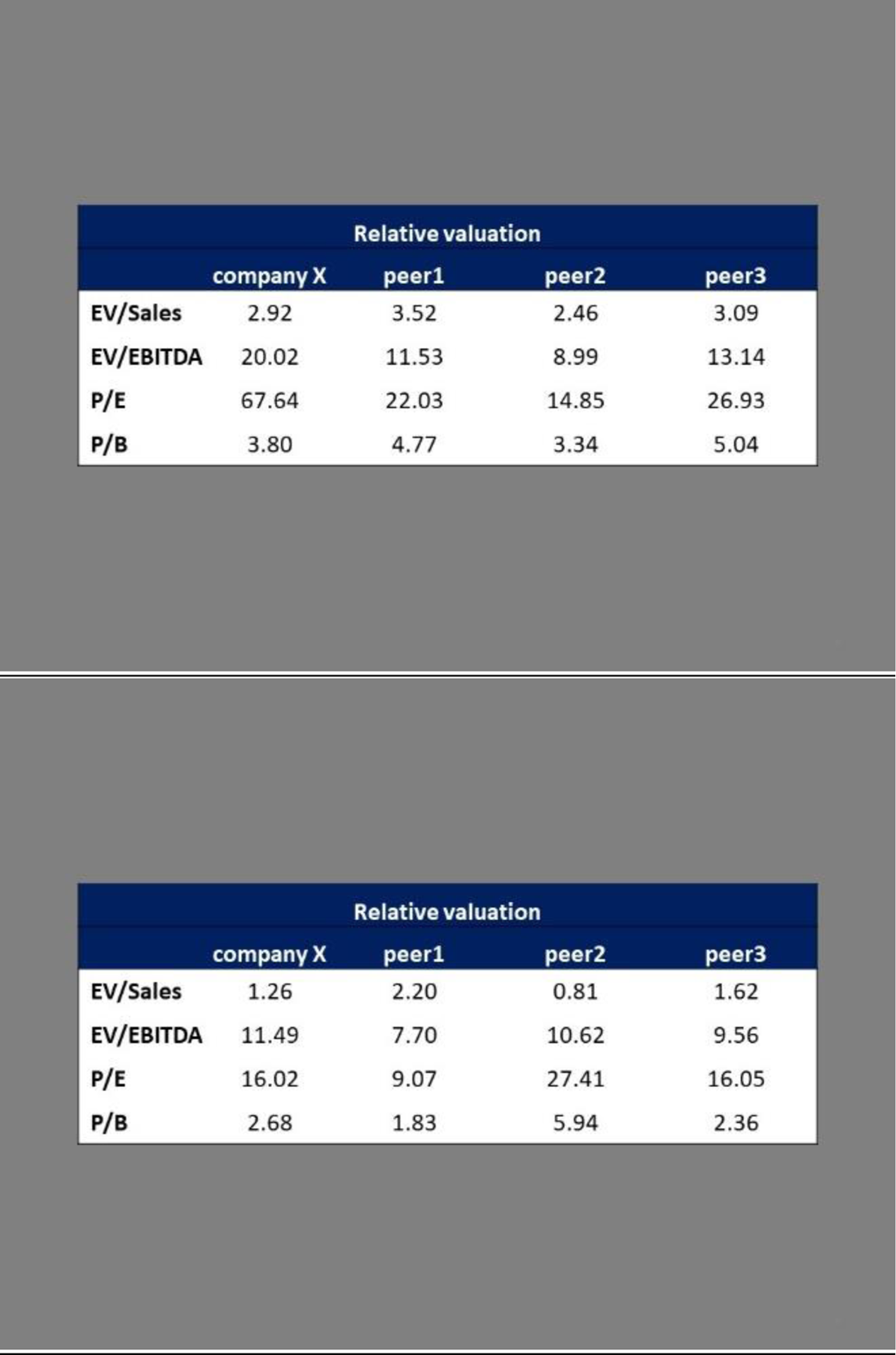
Two examples of the Fundamentals screen. The Fundamentals screen compared enterprise value (EV/Sales, EV/EBITDA) and price (P/E, P/B) metrics of the company to those of 3 peers.

**Fig. S5.**
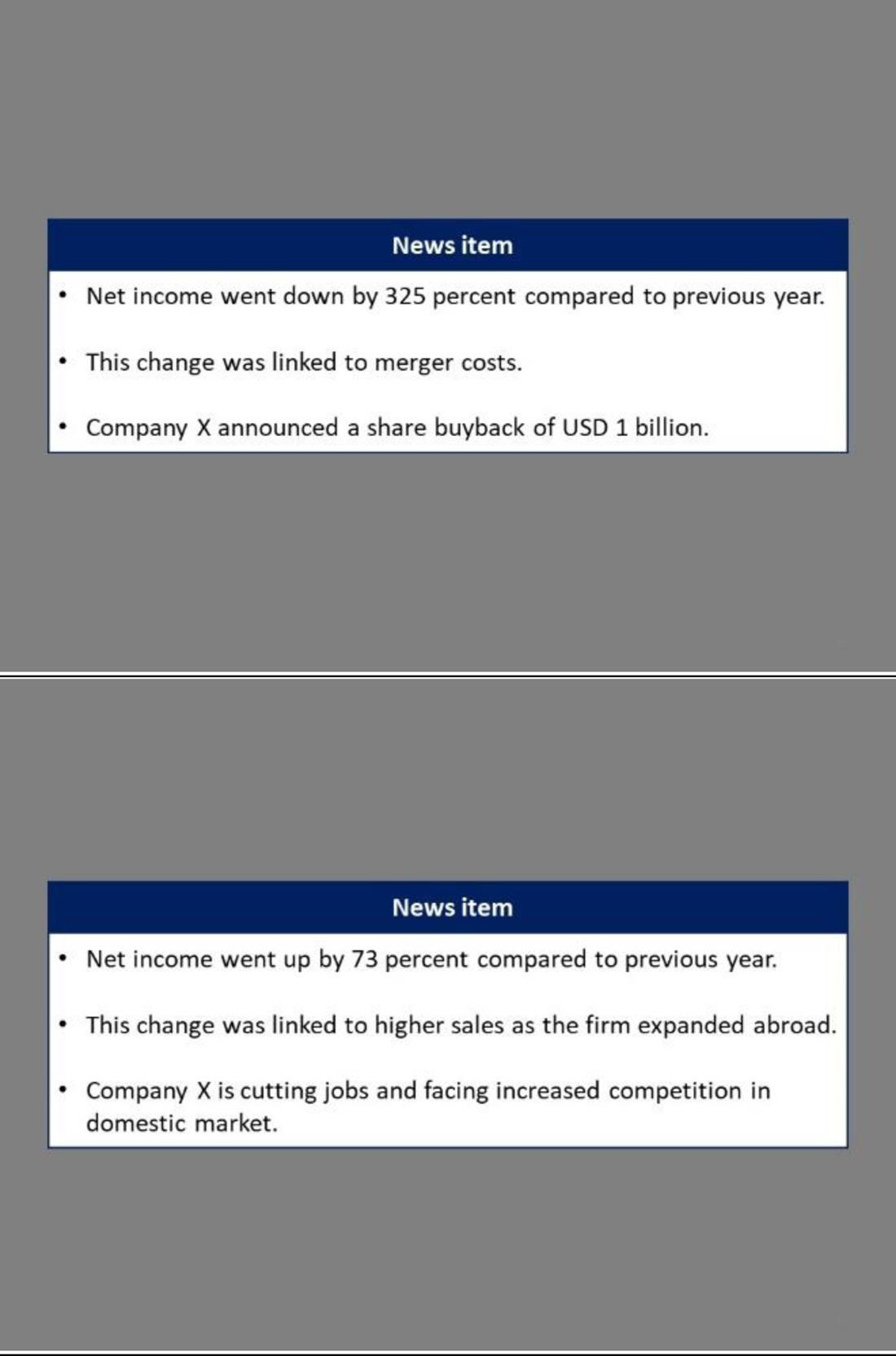
Two examples of the News Item screen. The News Item screen consisted of summaries of actual news, sampled from a Bloomberg terminal. Each news item consisted of three bullet points. The first bullet point always indicated the companies’ net income growth over the past year. The second bullet point provided an explanation for this (‘this change was related to…’) and the third item provided a prospect for the future.

## SI Appendix 2 fMRIprep preprocessing

Data was preprocessed using the standard pipeline of fMRIprep version 20.2.0 (27), based on Nipype (28). Specifically, anatomical T1-weighted (T1w) images were first corrected for intensity non-uniformity using N4BiasFieldCorrection (46), distributed with ANTs 2.3.3 (47). Next, T1w images were skull-stripped using antsBrainExtraction.sh workflow, followed by brain tissue segmentation of cerebrospinal fluid (CSF), white-matter (WM) and gray-matter (GM), using FAST on FSL v5.0.9 (48). Finally, T1w images were spatially normalized to the ICBM 152 Nonlinear Asymmetrical template 2009c (49). Functional images were preprocessed by first generating a reference volume and its skull-stripped version (standard fMRIprep). The BOLD reference was then co-registered to the T1w with a boundary-based registration cost-function (50), configured with nine degrees of freedom to account for remaining distortion in the BOLD reference. Next, head-motion parameters were estimated (transformation matrices and six corresponding rotation and translation parameters), before spatiotemporal filtering using MCFLIRT (51) and applying slice-time correction using 3dTshift from AFNI (52). BOLD time-series were then normalized to the same ICMB 152 Nonlinear space as the T1w images (49). Finally, global signal and framewise displacement (53) confounding time-series were calculated.

